# A step beyond: age and sex-dependent changes in multidimensional locomotor and gait patterns of voluntary motion of C57BL/6 mice

**DOI:** 10.64898/2026.10.26.761670

**Authors:** Husam Bensreti, Shawn Pavey, Billie Cullison, Kian Sheridan, Caihong Dai, Zihe Alex Zhang, Maribeth Johnson, David Maridas, Meghan E. McGee-Lawrence

**Affiliations:** Department of Cellular Biology and Anatomy, Medical College of Georgia, Augusta University, Augusta, GA; BlackBox Bio, Lewisville, TX; Department of Neuroscience & Regenerative Medicine, Medical College of, Georgia, Augusta University, Augusta, GA

**Keywords:** aging, gait, spontaneous activity, frailty, intrinsic capacity, vitality

## Abstract

Age-related decreases in physical activity drive frailty and reduced intrinsic capacity, promoting adverse health outcomes such as falls. Experimental tools for quantifying these metrics in preclinical models have lagged behind those used clinically; single-endpoint activity assessments (e.g., rotarod performance, treadmill activity) of rodents often require invasive handling and/or forced performance cues that may increase stress and variability, especially in aged animals. Here, we evaluated whether automated analysis of spontaneous movement using machine-learning-based software (Palmreader) could identify age-related changes in locomotor activity, gait, paw loading, or posture in male and female C57BL/6 mice across young, adult, and aged timepoints (4.5-, 7-, and 22 months of age). Parameters were quantified using the BlackBox One system during a 7-minute open-field recording. Principal component analysis demonstrated age-related separation of movement phenotypes, with individual parameters demonstrating increased hindlimb duty factor and longer stride lengths with age. Age-related changes in forelimb-to-hindlimb duty-factor imbalance, paw contact, and postural organization were also observed, and several gait alterations with age showed different patterns between males and females. Conventional measures of physical activity (e.g., distance traveled, walking time), in contrast, showed few age-related differences, particularly in females. These findings suggest that aging more consistently alters movement patterns rather than movement magnitude, and that gait and postural features may represent important age-related functional changes not captured by activity measurements alone. Automated assessment of spontaneous movement may therefore represent a useful complement to traditional physical activity testing, with the added benefit of minimal animal handling and short recording times.

## INTRODUCTION

Advances in medicine and improved living conditions have increased life expectancy worldwide [1]. As a result, the proportion of older adults in the population is growing and is expected to continue rising in the coming years. With this trajectory, the geriatric population is projected to exceed teenagers and young adults by 2040 and even surpass children by 2060 [2]. Because of this, it is important to develop new strategies aimed at ensuring that increased life expectancy will be accompanied by a good quality of life or healthspan; in the absence of such preservation, such as retained musculoskeletal mobility and physiological function, a longer lifespan may simply extend the period of poor health and disability that often occurs near end of life.

Frailty is correlated with loss of homeostasis and increased vulnerability to stressors, leading to an increased risk of disability, comorbidity, hospitalization, and death in the elderly [3]. This condition is defined based on the interaction of physiological, psychological, social, and environmental factors, representing the loss of physiological reserves across multiple organ systems [4]. Frailty is now recognized as a multifactorial syndrome characterized by a progressive decline in multiple organ systems, including the musculoskeletal, cardiovascular, metabolic, immune, and nervous systems. With the increase in life expectancy, frailty has become increasingly recognized as a rising health and socioeconomical problem, disproportionally affecting disadvantaged communities [5]. As a way to quantify frailty for cross-sectional and longitudinal assessments in human patients, a Frailty Index (FI) metric, also described as a Clinical Frailty Scale (CFS) in human patient populations, was developed by investigators to represent indices of several health deficits (e.g., clinical signs, symptoms, diseases, and disabilities) that accumulate with age [6–8]. This FI positively correlates with adverse outcomes, such as hospitalization [9], and represents a quantitative, multilevel individual evaluation that is considered a robust predictor for mortality [10]. While frailty has been in the scientific and clinical lexicon since at least the early 2000s, the more recent concept of ‘intrinsic capacity’ has emerged as an area of clinical interest in aging populations, representing the composite mental and physical internal attributes of an individual through measurements in the domains of locomotion, cognition, vitality, sensory function (vision and hearing), and psychological well-being or capacity [11–14]. These domain measurements of intrinsic capacity inversely correlate with adverse health outcomes such as fall risk and incident frailty [15].

Despite the substantial progress that has been made in identifying and characterizing frailty and intrinsic capacity in clinical settings, the experimental tools used to evaluate these metrics in preclinical models have lagged those used in human patient populations [11, 16, 17]. Several FI evaluation tools for rodents have been developed utilizing metrics of physical function and clinical presentation to measure frailty [18–24]. However, variability in measured endpoints among these FI methodologies is substantial, and many of the metrics rely on evaluations that require subjective assessments by a trained investigator [24–26]. To date, a unified approach for how to define and quantify frailty in pre-clinical models like mice has yet to be adopted by the field. Similarly, assessments of intrinsic capacity in rodents are still emerging. The recent HealthAge / INSPIRE study tracked intrinsic capacity-related measurements of vitality, locomotion, cognition, and psychological well-being in outbred male and female SWISS mice longitudinally and cross-sectionally between 6- and 24-months of age [27–29], but these results have yet to be replicated. Critically, many animal studies mainly rely on single-endpoint evaluations such as grip strength, rotarod performance, treadmill endurance, open field tests, or gait analysis as contributing variables to an overall FI or intrinsic capacity assessment [26, 27, 30]. While these tools have advanced the understanding of aging and musculoskeletal biology, they provide only a limited measurement of functional capacity when considered independently [30], and typically require animal handling by investigators and/or forced performance cues that may introduce stress and increase inter-operator variability, especially in aged animals. Importantly, these assays often provide little information on spontaneous physical activity, behavioral adaptability, fatigue, or recovery following physiological stress, all of which are increasingly recognized as major factors of frailty and age-related decline [31].

Consistent with this concept, recent clinical studies of human subjects demonstrated that continuous measurements of movement, mobility, and daily activity may identify decline in function earlier than traditional assessments of body composition or maximal strength alone [32]. This has supported increased interest in the development of continuous digital monitoring to record daily physical activity rather than relying on any single metric in controlled settings [33]. However, translation of this multi-factorial digital phenotyping framework into preclinical settings for aging and frailty research remains relatively limited, with most home-cage monitoring approaches focused on relatively macroscopic measures of activity, feeding, and social behavior rather than detailed measures of physical function [34, 35].

Next-generation instruments to address these shortfalls in assessment of voluntary activity and frailty-related measures in pre-clinical studies are being rapidly developed. As one such platform, the BlackBox One (and newer BlackBox Two) instrumentation produced by BlackBox Bio records detailed video of voluntary rodent movement and behavior while reducing background noise, animal handling, and variability in data collection. Similarly, the Palmreader software data management and analysis tool, which analyzes movement data collected on the BlackBox systems, is advertised as a machine learning-based platform that utilizes rodent behavioral analysis algorithms to automate measurement of paw luminance ratios (representing relative weight borne on each paw) and body pose estimation, enabling users to train convolutional neural networks to automate human-scored behaviors of interest. This platform has previously been utilized in pre-clinical rodent studies of pain-related alterations in gait and related behaviors [36, 37], functional deficits following spinal cord or nerve injuries [38, 39], and skeletal fractures in young mice [40], but had not yet been explored in the context of aging-related musculoskeletal decline and frailty. Here, we demonstrate that aging promotes a reduction in various activity-related parameters, as measured by the BlackBox One system in an unbiased and unstimulated manner, consistent with a deterioration in physical performance metrics indicative of musculoskeletal decline and frailty with age.

## MATERIALS AND METHODS

### Animal model

Male and female C57BL/6 mice at 7 months (young adult) and 22 months (aged adult) of age were obtained from the National Institute on Aging (NIA) aged rodent colony. An additional cohort of male and female C57BL/6 mice were bred in-house at Augusta University from breeding pairs obtained from The Jackson Laboratory and were studied at 4.5 months of age. Mice were group-housed in an accredited facility and maintained on a 12-hour light/dark cycle, and research was conducted according to guidelines provided by the National Institutes of Health (NIH) and under protocols approved by the Augusta University Institutional Animal Care and Use Committee (IACUC). Mice were maintained on a standard rodent diet (Teklad 2918, Envigo, Madison, WI, USA) and allowed food and water ad libitum.

### Open field locomotor assessment

Spontaneous continuous locomotor activity was recorded using the BlackBox One BB1-1R4 system on a 30cm x 30cm viewing area (BlackBox Biotech, TX, USA). IntellitouchO (FTIR) and TRANS voltage (brightness) values were set to 11.0 and 6.46 respectively. Mice were acclimated to the BlackBox chambers for 30 minutes on the day before recording. Before each recording session, the recording surface and animal chambers were cleaned. On the day of testing, mice were individually placed into the recording chamber, and spontaneous movement was recorded for 7 minutes under open-field (30 x30) area. Body mass was recorded on the same day. Analyses were performed using Palmreader software (v0.7.1), incorporating automated preprocessing and machine learning-based behavioral analysis. Automated analyses included measures of spontaneous locomotor activity (distance travelled, time spent near the center), gait (duty factors, stride lengths, temporal and spatial symmetry during detected gait bouts passing criteria such as paw alternation and smooth velocity), posture (stance width, joint angles and their variability), weightbearing (paw contact area, luminance, pressure index), and behavior (time spent rearing, grooming, turning) generated by the software. The full complement of measured parameters is shown in **Tables 1, 2, and 3**.

**Table 1.** Gait parameters generated by BlackBox analysis. Summary of gait-related parameters generated from automated analysis of spontaneous locomotion, including gait cycles and footsteps, paw-specific duty factor, limb phase, temporal and spatial symmetry/asymmetry, step width, stride length, velocity, and angular velocity. Paw-specific variables are reported for the left forepaw (LFP), right forepaw (RFP), left hindpaw (LHP), and right hindpaw (RHP). MAD, median absolute deviation. Variables reported as ratios, fractions, or phase measurements are unitless unless otherwise indicated. Definitions of software-specific gait parameters are provided according to the BlackBox output terminology

| **TABLE 1** |  |  |
| --- | --- | --- |
| **Parameter** | **Unit** | **Description** |
| Number_of_bouts | Count | Number of detected walking bouts. |
| Quadruped | Flag | Software quality flag readout; 1 if gait was quadrupedal, 0 if reverted to bipedal (e.g. hind limb paralysis, pipeline only outputs fore limb values) |
| Number_of_footsteps | Count | Number of detected paw steps. |
| Cycle | Count | Number of detected gait cycles (often approximately Number of bouts times 3) |
| Lfpaw_dorsal_steps | Count | Number of detected dorsal-surface contacts/steps by the left forepaw. |
| Lhpaw_dorsal_steps | Count | Number of detected dorsal-surface contacts/steps by the left hindpaw. |
| Rfpaw_dorsal_steps | Count | Number of detected dorsal-surface contacts/steps by the right forepaw. |
| Rhpaw_dorsal_steps | Count | Number of detected dorsal-surface contacts/steps by the right hindpaw. |
| Lfpaw_duty_factor | Ratio | Fraction of gait cycle spent in stance/contact for the left forepaw. |
| Lhpaw_duty_factor | Ratio | Fraction of gait cycle spent in stance/contact for the left hindpaw. |
| Rfpaw_duty_factor | Ratio | Fraction of gait cycle spent in stance/contact for the right forepaw. |
| Rhpaw_duty_factor | Ratio | Fraction of gait cycle spent in stance/contact for the right hindpaw. |
| Rfh_duty_factor_imbalance | Difference of Ratio | Difference in stance-time fraction between the limb pair indicated by the prefix (right side, fore minus hind) |
| R_duty_factor | Ratio | Fraction of gait cycle spent in stance/contact for the indicated limb pair (right side). |
| Lrh_duty_factor_imbalance | Difference of Ratio | Difference in stance-time fraction between the limb pair indicated by the prefix (left minus right, hind side). |
| H_duty_factor | Ratio | Fraction of gait cycle spent in stance/contact for the indicated limb pair (hind side). |
| Lrf_duty_factor_imbalance | Difference of | Difference in stance-time fraction between the limb pair indicated by |
|  | Ratio | the prefix (left minus right, fore side). |
| F_duty_factor | Ratio | Fraction of gait cycle spent in stance/contact for the indicated limb pair (fore side). |
| Lfh_duty_factor_imbalance | Difference of Ratio | Difference in stance-time fraction between the limb pair indicated by the prefix (left side, fore minus hind). |
| L_duty_factor | Ratio | Fraction of gait cycle spent in stance/contact for the indicated limb pair (left side). |
| Lfpaw_limb_phase | Ratio | Timing of left forepaw contact relative to the ipsilateral lhpaw within the gait cycle. |
| Lfpaw_limb_phase_mad | MAD of Ratio | Median absolute deviation of left forepaw limb phase across cycles. |
| Lfpaw_limb_phase_asymmetry | MAD of Ratio | Limb phase mad calculated within each cycle instead of full video. |
| Lhpaw_limb_phase | Ratio | Timing of left hindpaw contact relative to the ipsilateral lfpaw within the gait cycle. |
| Lhpaw_limb_phase_mad | MAD of Ratio | Median absolute deviation of left hindpaw limb phase across cycles. |
| Lhpaw_limb_phase_asymmetry | MAD of Ratio | Limb phase mad calculated within each cycle instead of full video. |
| Rfpaw_limb_phase | Ratio | Timing of right forepaw contact relative to the ipsilateral rhpaw within the gait cycle. |
| Rfpaw_limb_phase_mad | MAD of Ratio | Median absolute deviation of right forepaw limb phase across cycles. |
| Rfpaw_limb_phase_asymmetry | MAD of Ratio | Limb phase mad calculated within each cycle instead of full video. |
| Rhpaw_limb_phase | Ratio | Timing of right hindpaw contact relative to the ipsilateral rfpaw within the gait cycle. |
| Rhpaw_limb_phase_mad | MAD of Ratio | Median absolute deviation of right hindpaw limb phase across cycles. |
| Rhpaw_limb_phase_asymmetry | MAD of Ratio | Limb phase mad calculated within each cycle instead of full video. |
| Lfpaw_temporal_symmetry | Ratio | Ratio of contralateral to current footstrike time divided by stride time of the contralateral (0.5 is perfectly symmetrical, > 0.5 is delayed) |
| Lfpaw_temporal_asymmetry | Difference of Ratio | Average of absolute difference from perfectly symmetrical gait (0.5) |
| Lhpaw_temporal_symmetry | Ratio | Ratio of contralateral to current footstrike time divided by stride time of the contralateral (0.5 is perfectly symmetrical, > 0.5 is delayed) |
| Lhpaw_temporal_asymmetry | Difference of Ratio | Average of absolute difference from perfectly symmetrical gait (0.5) |
| Rfpaw_temporal_symmetry | Ratio | Ratio of contralateral to current footstrike time divided by stride time of the contralateral (0.5 is perfectly symmetrical, > 0.5 is delayed) |
| Rfpaw_temporal_asymmetry | Difference of Ratio | Average of absolute difference from perfectly symmetrical gait (0.5) |
| Rhpaw_temporal_symmetry | Ratio | Ratio of contralateral to current footstrike time divided by stride time of the contralateral (0.5 is perfectly symmetrical, > 0.5 is delayed) |
| Rhpaw_temporal_asymmetry | Difference of Ratio | Average of absolute difference from perfectly symmetrical gait (0.5) |
| Lfpaw_step_width | M | Lateral spacing of left forepaw step relative to the contralateral. |
| Lfpaw_stride_length | M | Distance between successive placements of the left forepaw. |
| Lfpaw_spatial_symmetry | Ratio | Ratio of perpendicular distance between previous contralateral and current over contralateral stride length (0.5 = symmetrical placement) |
| Lfpaw_spatial_asymmetry | Difference of | Average of absolute difference from perfectly symmetrical gait (0.5) |
|  | Ratio |  |
| Lhpaw_step_width | M | Lateral spacing of left hindpaw step relative to the contralateral. |
| Lhpaw_stride_length | M | Distance between successive placements of the left forepaw. |
| Lhpaw_spatial_symmetry | Ratio | Ratio of perpendicular distance between previous contralateral and current over contralateral stride length (0.5 = symmetrical placement) |
| Lhpaw_spatial_asymmetry | Difference of Ratio | Average of absolute difference from perfectly symmetrical gait (0.5) |
| Rfpaw_step_width | M | Lateral spacing of right forepaw step relative to the contralateral. |
| Rfpaw_stride_length | M | Distance between successive placements of the left forepaw. |
| Rfpaw_spatial_symmetry | Ratio | Ratio of perpendicular distance between previous contralateral and current over contralateral stride length (0.5 = symmetrical placement) |
| Rfpaw_spatial_asymmetry | Difference of Ratio | Average of absolute difference from perfectly symmetrical gait (0.5) |
| Rhpaw_step_width | M | Lateral spacing of right hindpaw step relative to the contralateral/locomotor reference. |
| Rhpaw_stride_length | M | Distance between successive placements of the left forepaw. |
| Rhpaw_spatial_symmetry | Ratio | Ratio of perpendicular distance between previous contralateral and current over contralateral stride length (0.5 = symmetrical placement) |
| Rhpaw_spatial_asymmetry | Difference of Ratio | Average of absolute difference from perfectly symmetrical gait (0.5) |
| Velocity | m/s | Mean movement speed. |
| Angular_velocity | rad/s | Rate of change in heading during locomotion. |

**Table 2.**
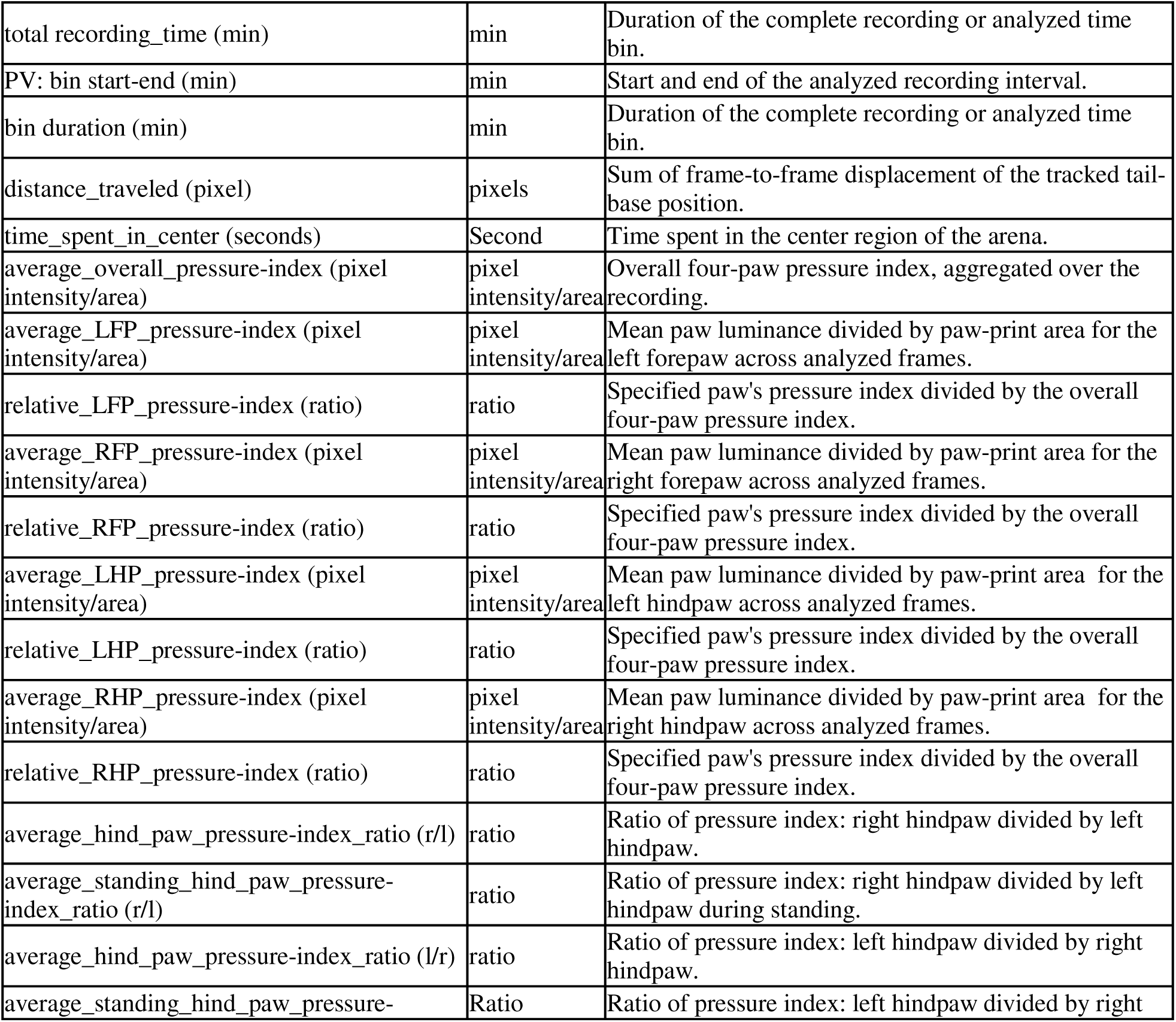
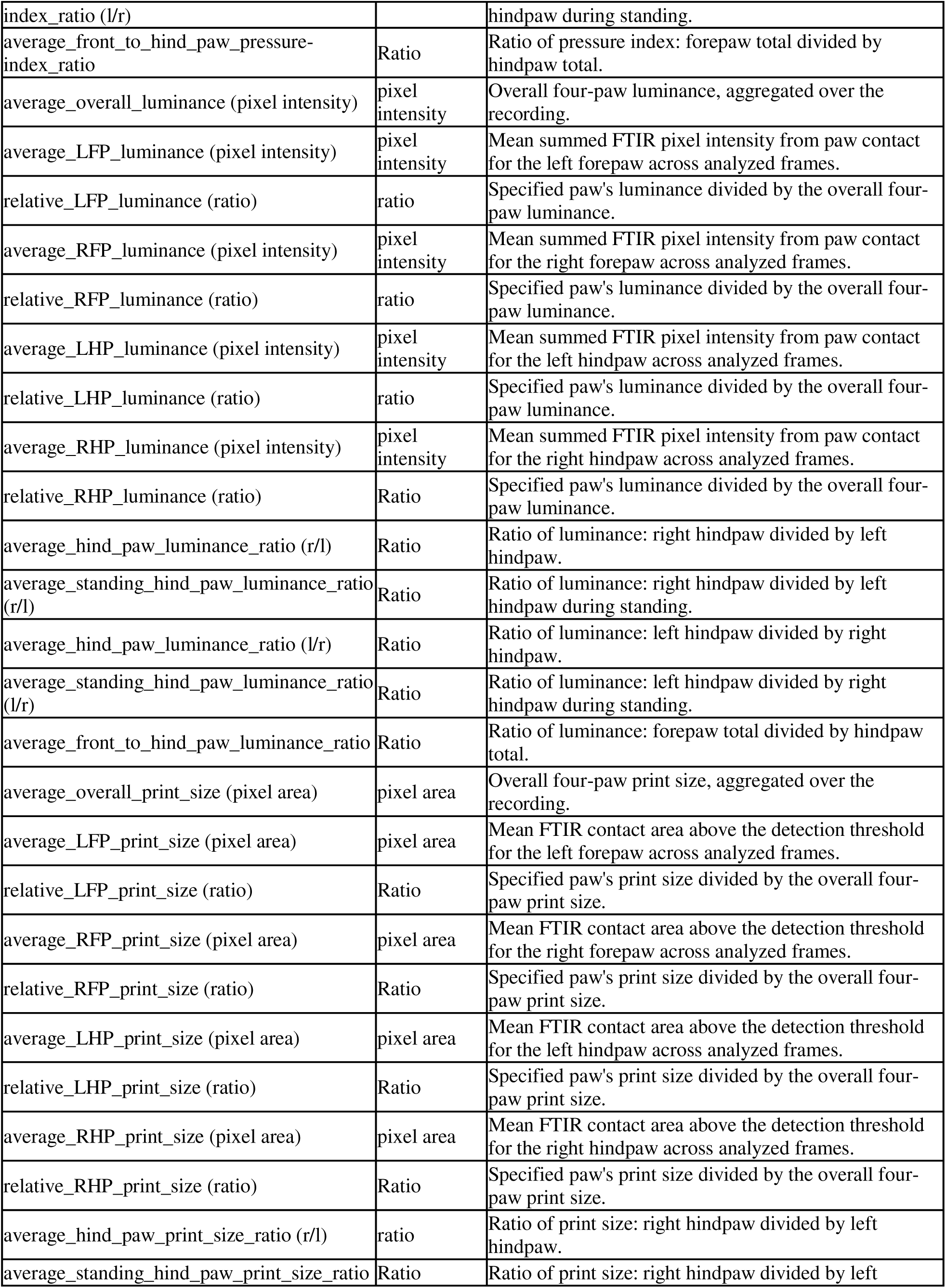
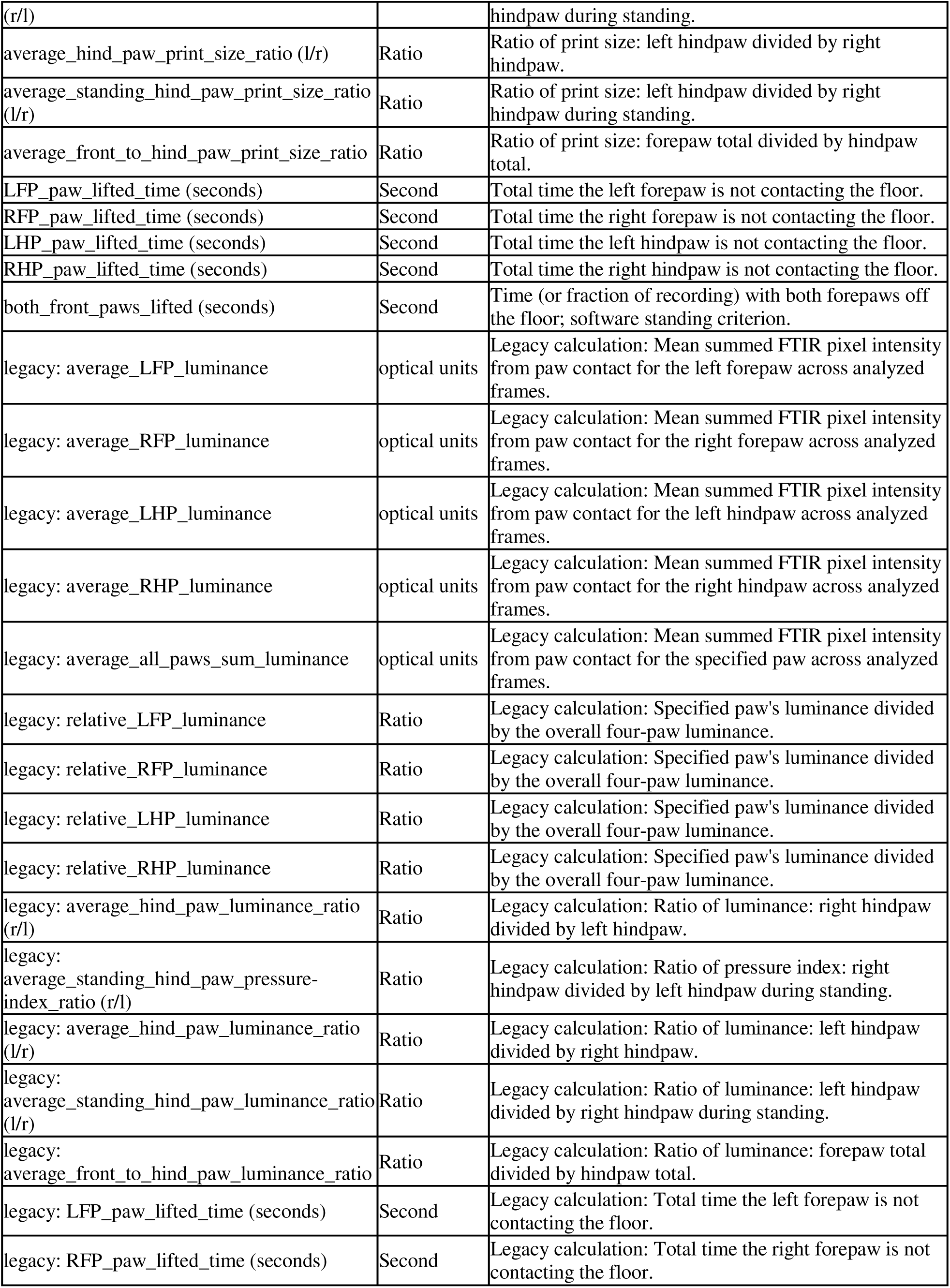
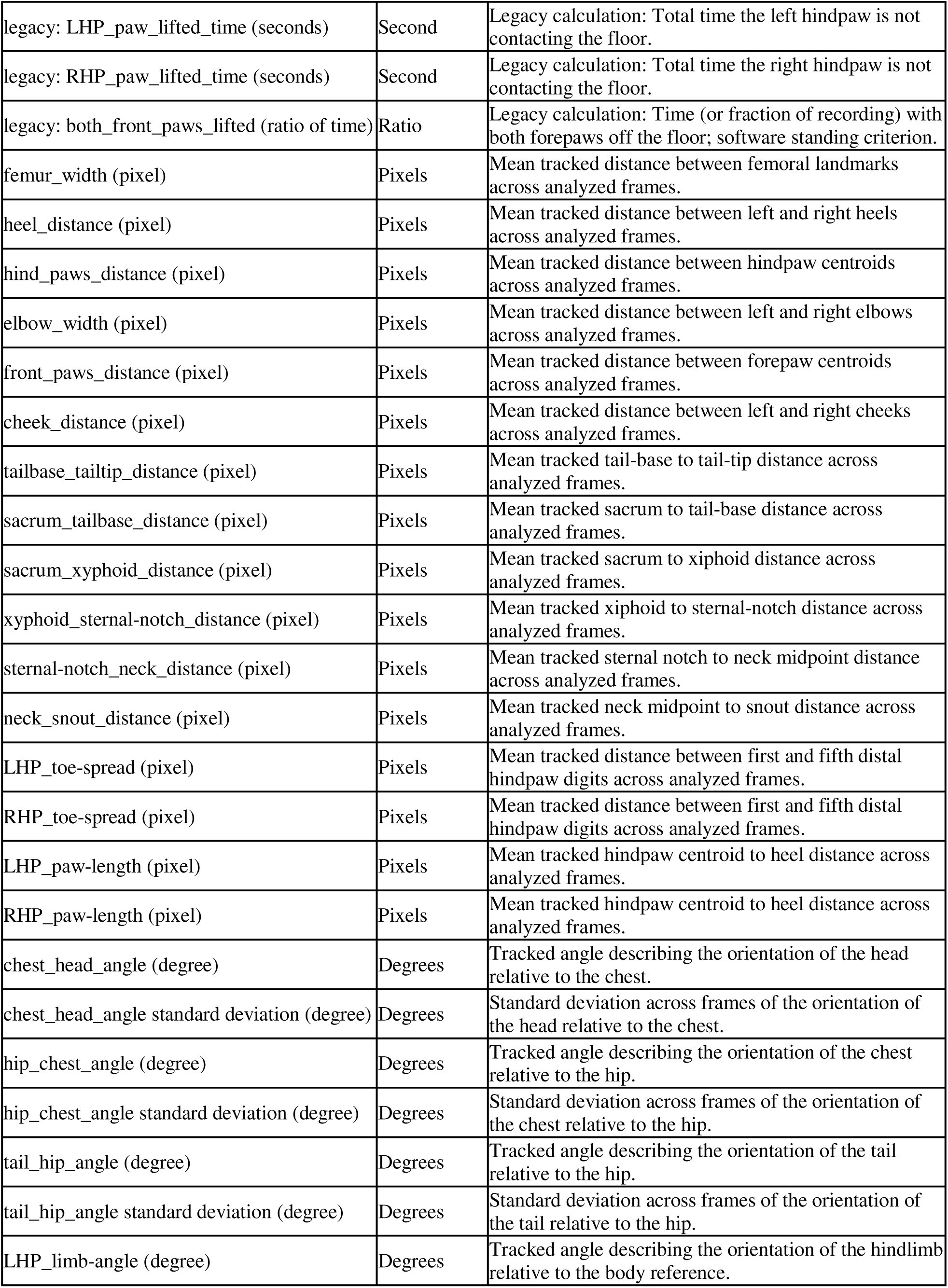
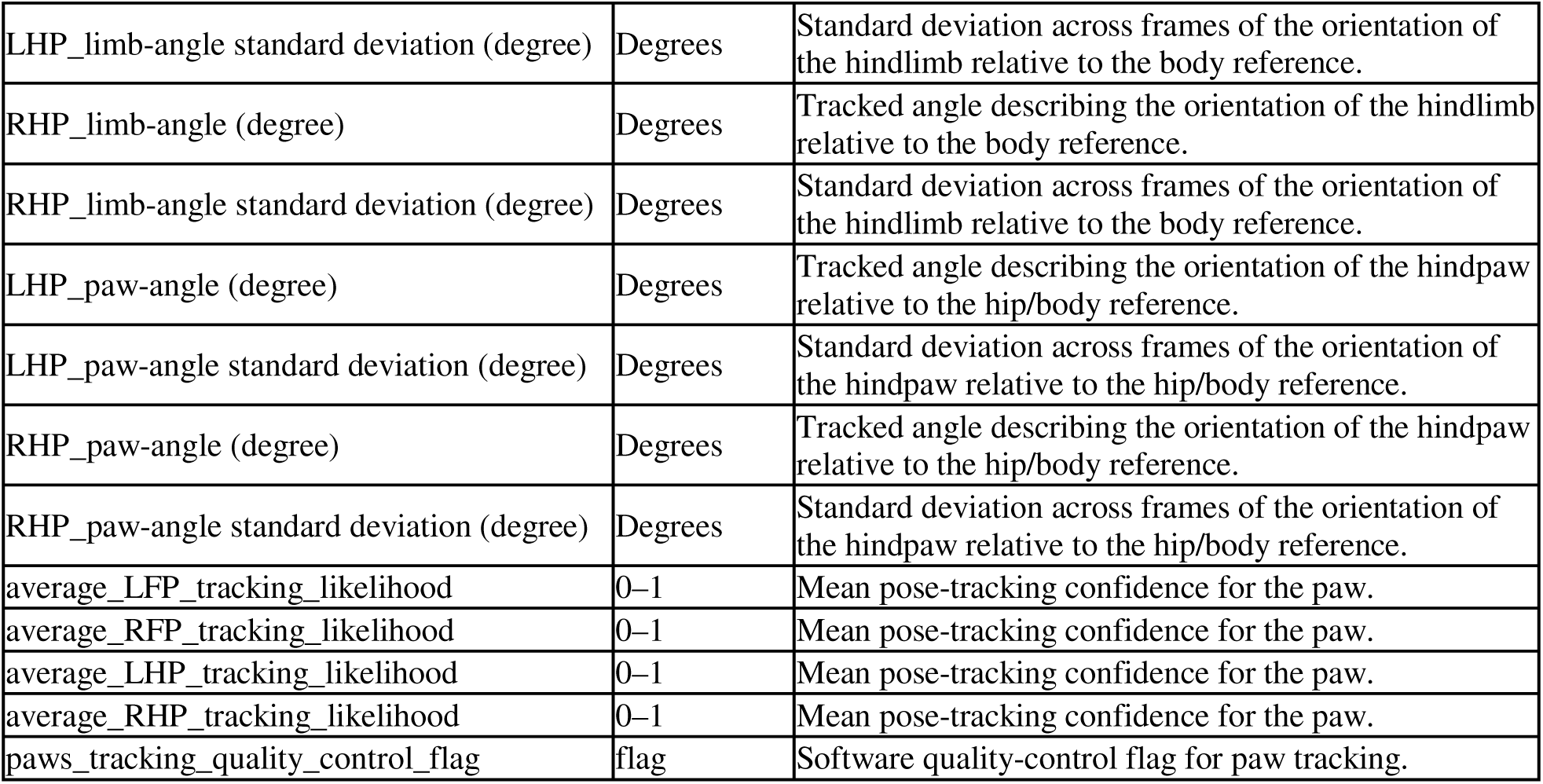
General activity, plantar-contact, and postural parameters generated by BlackBox analysis. Summary of general activity, paw-contact/loading, and postural measurements obtained during spontaneous activity. General measures include recording duration, distance traveled, and time spent in the center. Plantar measurements include paw-print size, luminance, optical pressure index, relative paw measurements, forepaw-to-hindpaw and left-to-right ratios, and paw-lifted time. Postural measurements include distances and angles between tracked anatomical landmarks, hindpaw dimensions, and variability in body angles.

| **TABLE 2** |  |  |
| --- | --- | --- |
| Parameter | Unit | Description |
| total recording_time (min) | min | Duration of the complete recording or analyzed time bin. |
| PV: bin start-end (min) | min | Start and end of the analyzed recording interval. |
| bin duration (min) | min | Duration of the complete recording or analyzed time bin. |
| distance_traveled (pixel) | pixels | Sum of frame-to-frame displacement of the tracked tail-base position. |
| time_spent_in_center (seconds) | Second | Time spent in the center region of the arena. |
| average_overall_pressure-index (pixel intensity/area) | pixel intensity/area | Overall four-paw pressure index, aggregated over the recording. |
| average_LFP_pressure-index (pixel intensity/area) | pixel intensity/area | Mean paw luminance divided by paw-print area for the left forepaw across analyzed frames. |
| relative_LFP_pressure-index (ratio) | ratio | Specified paw's pressure index divided by the overall four-paw pressure index. |
| average_RFP_pressure-index (pixel intensity/area) | pixel intensity/area | Mean paw luminance divided by paw-print area for the right forepaw across analyzed frames. |
| relative_RFP_pressure-index (ratio) | ratio | Specified paw's pressure index divided by the overall four-paw pressure index. |
| average_LHP_pressure-index (pixel intensity/area) | pixel intensity/area | Mean paw luminance divided by paw-print area for the left hindpaw across analyzed frames. |
| relative_LHP_pressure-index (ratio) | ratio | Specified paw's pressure index divided by the overall four-paw pressure index. |
| average_RHP_pressure-index (pixel intensity/area) | pixel intensity/area | Mean paw luminance divided by paw-print area for the right hindpaw across analyzed frames. |
| relative_RHP_pressure-index (ratio) | ratio | Specified paw's pressure index divided by the overall four-paw pressure index. |
| average_hind_paw_pressure-index_ratio (r/l) | ratio | Ratio of pressure index: right hindpaw divided by left hindpaw. |
| average_standing_hind_paw_pressure-index_ratio (r/l) | ratio | Ratio of pressure index: right hindpaw divided by left hindpaw during standing. |
| average_hind_paw_pressure-index_ratio (l/r) | ratio | Ratio of pressure index: left hindpaw divided by right hindpaw. |
| average_standing_hind_paw_pressure- | Ratio | Ratio of pressure index: left hindpaw divided by right |
| index_ratio (l/r) |  | hindpaw during standing. |
| average_front_to_hind_paw_pressure-index_ratio | Ratio | Ratio of pressure index: forepaw total divided by hindpaw total. |
| average_overall_luminance (pixel intensity) | pixel intensity | Overall four-paw luminance, aggregated over the recording. |
| average_LFP_luminance (pixel intensity) | pixel intensity | Mean summed FTIR pixel intensity from paw contact for the left forepaw across analyzed frames. |
| relative_LFP_luminance (ratio) | ratio | Specified paw's luminance divided by the overall four-paw luminance. |
| average_RFP_luminance (pixel intensity) | pixel intensity | Mean summed FTIR pixel intensity from paw contact for the right forepaw across analyzed frames. |
| relative_RFP_luminance (ratio) | ratio | Specified paw's luminance divided by the overall four-paw luminance. |
| average_LHP_luminance (pixel intensity) | pixel intensity | Mean summed FTIR pixel intensity from paw contact for the left hindpaw across analyzed frames. |
| relative_LHP_luminance (ratio) | ratio | Specified paw's luminance divided by the overall four-paw luminance. |
| average_RHP_luminance (pixel intensity) | pixel intensity | Mean summed FTIR pixel intensity from paw contact for the right hindpaw across analyzed frames. |
| relative_RHP_luminance (ratio) | Ratio | Specified paw's luminance divided by the overall four-paw luminance. |
| average_hind_paw_luminance_ratio (r/l) | Ratio | Ratio of luminance: right hindpaw divided by left hindpaw. |
| average_standing_hind_paw_luminance_ratio (r/l) | Ratio | Ratio of luminance: right hindpaw divided by left hindpaw during standing. |
| average_hind_paw_luminance_ratio (l/r) | Ratio | Ratio of luminance: left hindpaw divided by right hindpaw. |
| average_standing_hind_paw_luminance_ratio (l/r) | Ratio | Ratio of luminance: left hindpaw divided by right hindpaw during standing. |
| average_front_to_hind_paw_luminance_ratio | Ratio | Ratio of luminance: forepaw total divided by hindpaw total. |
| average_overall_print_size (pixel area) | pixel area | Overall four-paw print size, aggregated over the recording. |
| average_LFP_print_size (pixel area) | pixel area | Mean FTIR contact area above the detection threshold for the left forepaw across analyzed frames. |
| relative_LFP_print_size (ratio) | Ratio | Specified paw's print size divided by the overall four-paw print size. |
| average_RFP_print_size (pixel area) | pixel area | Mean FTIR contact area above the detection threshold for the right forepaw across analyzed frames. |
| relative_RFP_print_size (ratio) | Ratio | Specified paw's print size divided by the overall four-paw print size. |
| average_LHP_print_size (pixel area) | pixel area | Mean FTIR contact area above the detection threshold for the left hindpaw across analyzed frames. |
| relative_LHP_print_size (ratio) | Ratio | Specified paw's print size divided by the overall four-paw print size. |
| average_RHP_print_size (pixel area) | pixel area | Mean FTIR contact area above the detection threshold for the right hindpaw across analyzed frames. |
| relative_RHP_print_size (ratio) | Ratio | Specified paw's print size divided by the overall four-paw print size. |
| average_hind_paw_print_size_ratio (r/l) | ratio | Ratio of print size: right hindpaw divided by left hindpaw. |
| average_standing_hind_paw_print_size_ratio | Ratio | Ratio of print size: right hindpaw divided by left |
| (r/l) |  | hindpaw during standing. |
| average_hind_paw_print_size_ratio (l/r) | Ratio | Ratio of print size: left hindpaw divided by right hindpaw. |
| average_standing_hind_paw_print_size_ratio (l/r) | Ratio | Ratio of print size: left hindpaw divided by right hindpaw during standing. |
| average_front_to_hind_paw_print_size_ratio | Ratio | Ratio of print size: forepaw total divided by hindpaw total. |
| LFP_paw_lifted_time (seconds) | Second | Total time the left forepaw is not contacting the floor. |
| RFP_paw_lifted_time (seconds) | Second | Total time the right forepaw is not contacting the floor. |
| LHP_paw_lifted_time (seconds) | Second | Total time the left hindpaw is not contacting the floor. |
| RHP_paw_lifted_time (seconds) | Second | Total time the right hindpaw is not contacting the floor. |
| both_front_paws_lifted (seconds) | Second | Time (or fraction of recording) with both forepaws off the floor; software standing criterion. |
| legacy: average_LFP_luminance | optical units | Legacy calculation: Mean summed FTIR pixel intensity from paw contact for the left forepaw across analyzed frames. |
| legacy: average_RFP_luminance | optical units | Legacy calculation: Mean summed FTIR pixel intensity from paw contact for the right forepaw across analyzed frames. |
| legacy: average_LHP_luminance | optical units | Legacy calculation: Mean summed FTIR pixel intensity from paw contact for the left hindpaw across analyzed frames. |
| legacy: average_RHP_luminance | optical units | Legacy calculation: Mean summed FTIR pixel intensity from paw contact for the right hindpaw across analyzed frames. |
| legacy: average_all_paws_sum_luminance | optical units | Legacy calculation: Mean summed FTIR pixel intensity from paw contact for the specified paw across analyzed frames. |
| legacy: relative_LFP_luminance | Ratio | Legacy calculation: Specified paw's luminance divided by the overall four-paw luminance. |
| legacy: relative_RFP_luminance | Ratio | Legacy calculation: Specified paw's luminance divided by the overall four-paw luminance. |
| legacy: relative_LHP_luminance | Ratio | Legacy calculation: Specified paw's luminance divided by the overall four-paw luminance. |
| legacy: relative_RHP_luminance | Ratio | Legacy calculation: Specified paw's luminance divided by the overall four-paw luminance. |
| legacy: average_hind_paw_luminance_ratio (r/l) | Ratio | Legacy calculation: Ratio of luminance: right hindpaw divided by left hindpaw. |
| legacy: average_standing_hind_paw_pressure-index_ratio (r/l) | Ratio | Legacy calculation: Ratio of pressure index: right hindpaw divided by left hindpaw during standing. |
| legacy: average_hind_paw_luminance_ratio (l/r) | Ratio | Legacy calculation: Ratio of luminance: left hindpaw divided by right hindpaw. |
| legacy: average_standing_hind_paw_luminance_ratio (l/r) | Ratio | Legacy calculation: Ratio of luminance: left hindpaw divided by right hindpaw during standing. |
| legacy: average_front_to_hind_paw_luminance_ratio | Ratio | Legacy calculation: Ratio of luminance: forepaw total divided by hindpaw total. |
| legacy: LFP_paw_lifted_time (seconds) | Second | Legacy calculation: Total time the left forepaw is not contacting the floor. |
| legacy: RFP_paw_lifted_time (seconds) | Second | Legacy calculation: Total time the right forepaw is not contacting the floor. |
| legacy: LHP_paw_lifted_time (seconds) | Second | Legacy calculation: Total time the left hindpaw is not contacting the floor. |
| legacy: RHP_paw_lifted_time (seconds) | Second | Legacy calculation: Total time the right hindpaw is not contacting the floor. |
| legacy: both_front_paws_lifted (ratio of time) | Ratio | Legacy calculation: Time (or fraction of recording) with both forepaws off the floor; software standing criterion. |
| femur_width (pixel) | Pixels | Mean tracked distance between femoral landmarks across analyzed frames. |
| heel_distance (pixel) | Pixels | Mean tracked distance between left and right heels across analyzed frames. |
| hind_paws_distance (pixel) | Pixels | Mean tracked distance between hindpaw centroids across analyzed frames. |
| elbow_width (pixel) | Pixels | Mean tracked distance between left and right elbows across analyzed frames. |
| front_paws_distance (pixel) | Pixels | Mean tracked distance between forepaw centroids across analyzed frames. |
| cheek_distance (pixel) | Pixels | Mean tracked distance between left and right cheeks across analyzed frames. |
| tailbase_tailtip_distance (pixel) | Pixels | Mean tracked tail-base to tail-tip distance across analyzed frames. |
| sacrum_tailbase_distance (pixel) | Pixels | Mean tracked sacrum to tail-base distance across analyzed frames. |
| sacrum_xyphoid_distance (pixel) | Pixels | Mean tracked sacrum to xiphoid distance across analyzed frames. |
| xyphoid_sternal-notch_distance (pixel) | Pixels | Mean tracked xiphoid to sternal-notch distance across analyzed frames. |
| sternal-notch_neck_distance (pixel) | Pixels | Mean tracked sternal notch to neck midpoint distance across analyzed frames. |
| neck_snout_distance (pixel) | Pixels | Mean tracked neck midpoint to snout distance across analyzed frames. |
| LHP_toe-spread (pixel) | Pixels | Mean tracked distance between first and fifth distal hindpaw digits across analyzed frames. |
| RHP_toe-spread (pixel) | Pixels | Mean tracked distance between first and fifth distal hindpaw digits across analyzed frames. |
| LHP_paw-length (pixel) | Pixels | Mean tracked hindpaw centroid to heel distance across analyzed frames. |
| RHP_paw-length (pixel) | Pixels | Mean tracked hindpaw centroid to heel distance across analyzed frames. |
| chest_head_angle (degree) | Degrees | Tracked angle describing the orientation of the head relative to the chest. |
| chest_head_angle standard deviation (degree) | Degrees | Standard deviation across frames of the orientation of the head relative to the chest. |
| hip_chest_angle (degree) | Degrees | Tracked angle describing the orientation of the chest relative to the hip. |
| hip_chest_angle standard deviation (degree) | Degrees | Standard deviation across frames of the orientation of the chest relative to the hip. |
| tail_hip_angle (degree) | Degrees | Tracked angle describing the orientation of the tail relative to the hip. |
| tail_hip_angle standard deviation (degree) | Degrees | Standard deviation across frames of the orientation of the tail relative to the hip. |
| LHP_limb-angle (degree) | Degrees | Tracked angle describing the orientation of the hindlimb relative to the body reference. |
| LHP_limb-angle standard deviation (degree) | Degrees | Standard deviation across frames of the orientation of the hindlimb relative to the body reference. |
| RHP_limb-angle (degree) | Degrees | Tracked angle describing the orientation of the hindlimb relative to the body reference. |
| RHP_limb-angle standard deviation (degree) | Degrees | Standard deviation across frames of the orientation of the hindlimb relative to the body reference. |
| LHP_paw-angle (degree) | Degrees | Tracked angle describing the orientation of the hindpaw relative to the hip/body reference. |
| LHP_paw-angle standard deviation (degree) | Degrees | Standard deviation across frames of the orientation of the hindpaw relative to the hip/body reference. |
| RHP_paw-angle (degree) | Degrees | Tracked angle describing the orientation of the hindpaw relative to the hip/body reference. |
| RHP_paw-angle standard deviation (degree) | Degrees | Standard deviation across frames of the orientation of the hindpaw relative to the hip/body reference. |
| average_LFP_tracking_likelihood | 0–1 | Mean pose-tracking confidence for the paw. |
| average_RFP_tracking_likelihood | 0–1 | Mean pose-tracking confidence for the paw. |
| average_LHP_tracking_likelihood | 0–1 | Mean pose-tracking confidence for the paw. |
| average_RHP_tracking_likelihood | 0–1 | Mean pose-tracking confidence for the paw. |
| paws_tracking_quality_control_flag | flag | Software quality-control flag for paw tracking. |

**Table 3.**
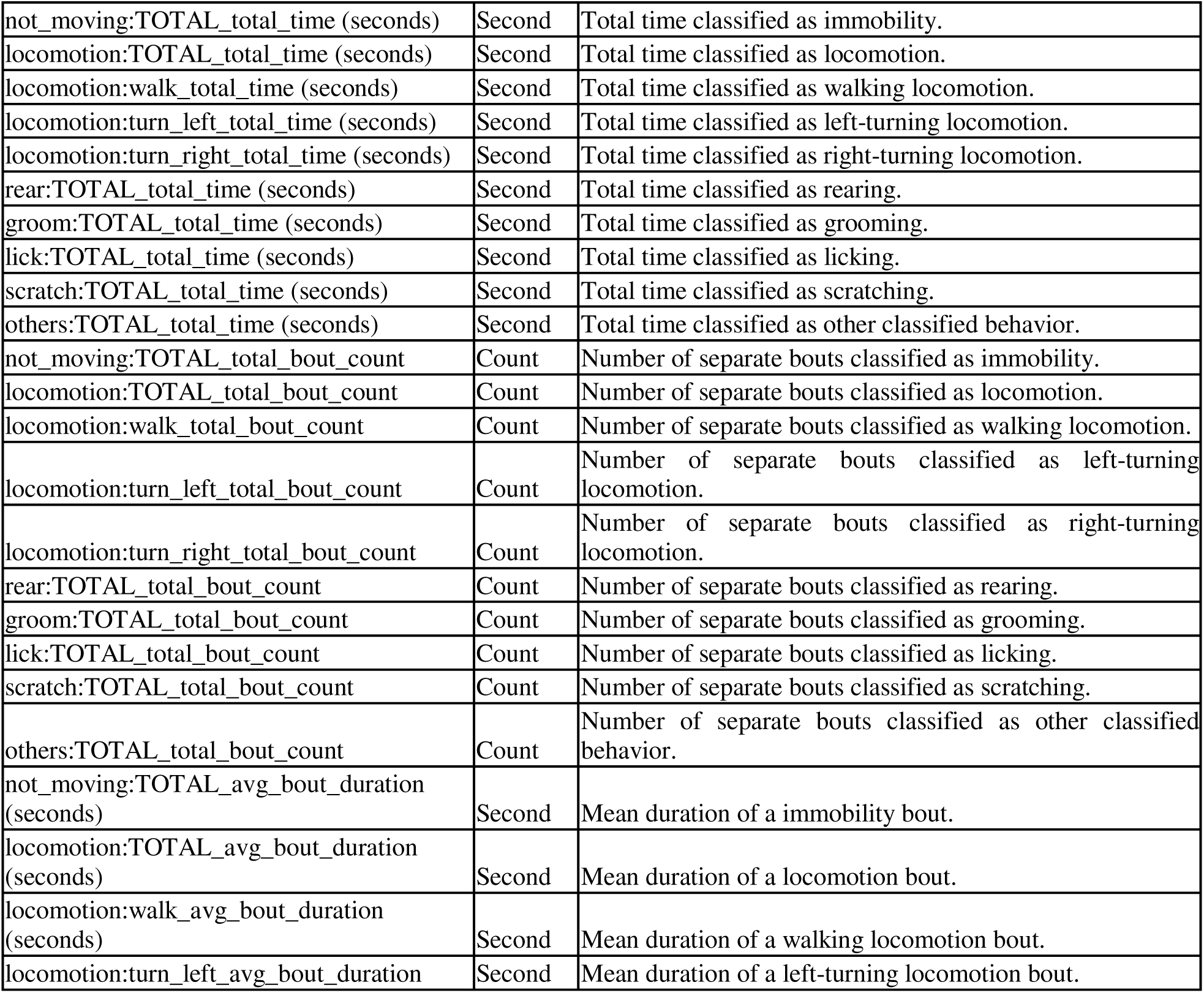
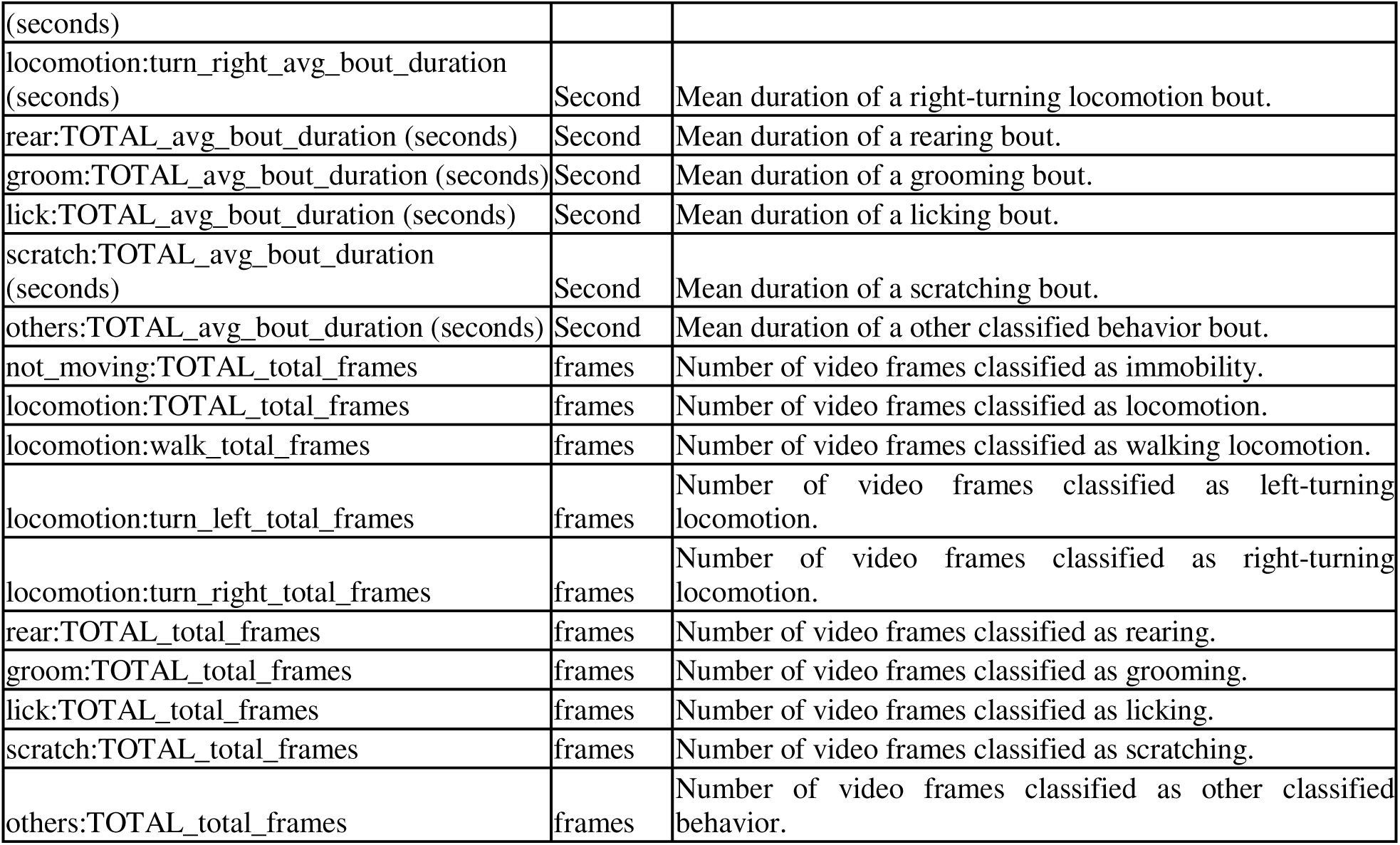
Ethogram parameters generated by BlackBox analysis. Summary of automatically classified behavioral measures during spontaneous activity. Behavioral categories include immobility (not moving), locomotion, walking, left and right turning, rearing, grooming, licking, scratching, and other classified behaviors. For each behavior, outputs include total time, total number of bouts, average bout duration, and total number of video frames assigned to that behavior. Time variables are reported in seconds, bout counts represent the number of discrete behavioral episodes, and frame counts represent the number of video frames classified within each behavioral category.

### Statistical analysis

Data were modeled using ordinary least squares, with sex and age as fixed effects. The model included normalizing z-scored covariates, which were also applied to per subject data points on all plots. These covariates were body mass (for postural, weightbearing, gait features) and both linear and angular velocity (gait, especially crucial to interpretations of symmetry and duty factors) [41], with no covariates applied to automated behavior scoring (rearing detection). Principal Component Analysis (PCA) data were also standardized (zero mean, unit variance) before fitting, with separate fits by sex so principal component loadings revealed age-related progression within each sex. The model’s estimated marginal means and their 95% confidence intervals were plotted over the individual data points for box and PCA plots. Confidence intervals and p-values were based on the t distribution using the model’s residual degrees of freedom. For groupwise comparisons of rearing behavior, posture, weightbearing, and gait, data were plotted for males and females separately, which reflects the structure of the 6 comparisons (3 per sex) included in the application of the Benjamini-Hochberg false discovery rate procedure. Exploratory feature heatmaps show z-scored estimated marginal means with covariate correction for pre-selected/engineered feature candidates, and a Bonferroni correction across rows.

For all reported metrics, statistical analyses consider all three groups (4.5 months old, 7 months old, and 22 months old) simultaneously, recognizing the limitation that the 4.5-month-old animals were obtained from a different rodent stock than the older groups. Subsequent comparisons featuring only the 7-month-old vs. 22-month-old mice, where animals were obtained from the same source colony, are highlighted where appropriate.

## RESULTS

### Aging produces distinct multivariate locomotor and gait characteristics

To determine whether spontaneous movement patterns change with age, male and female C57BL/6 mice at 4.5, 7, and 22 months of age were evaluated using the BlackBox One system to quantify measurements of locomotor activity, gait, paw placement and loading, and posture. Since many of these measurements are related, principal component analysis (PCA) was used to examine how the overall movement phenotype differed across age groups.

When all measured parameters were included, age-related differences were observed in both females and males (**Figure 1A-B**). The separation across principal component 1 (PC1) distinguished 4.5- and 7-month-old groups in both sexes, characterized by a shift towards the hind paws in terms of weightbearing. The separation across PC2 distinguished 7- and 22-month-old groups in both cases. In both sexes, this progression was characterized by increased stride length and stance width, with reduced locomotion, rearing (or fore paw lifting) and gait/postural variability. In males this affected a broader range of postural features, such as *Fh Paw Distance Ratio* (the ratio of fore paw distance to hind paw). Overall, the unsupervised PCA showed orthogonal shifts from 4.5 months to 7 months versus 7 months to 22 months, suggesting that the latter was not simply a continuation of skeletal maturation but rather driven by a separate set of phenotypic changes.

**Figure 1.**
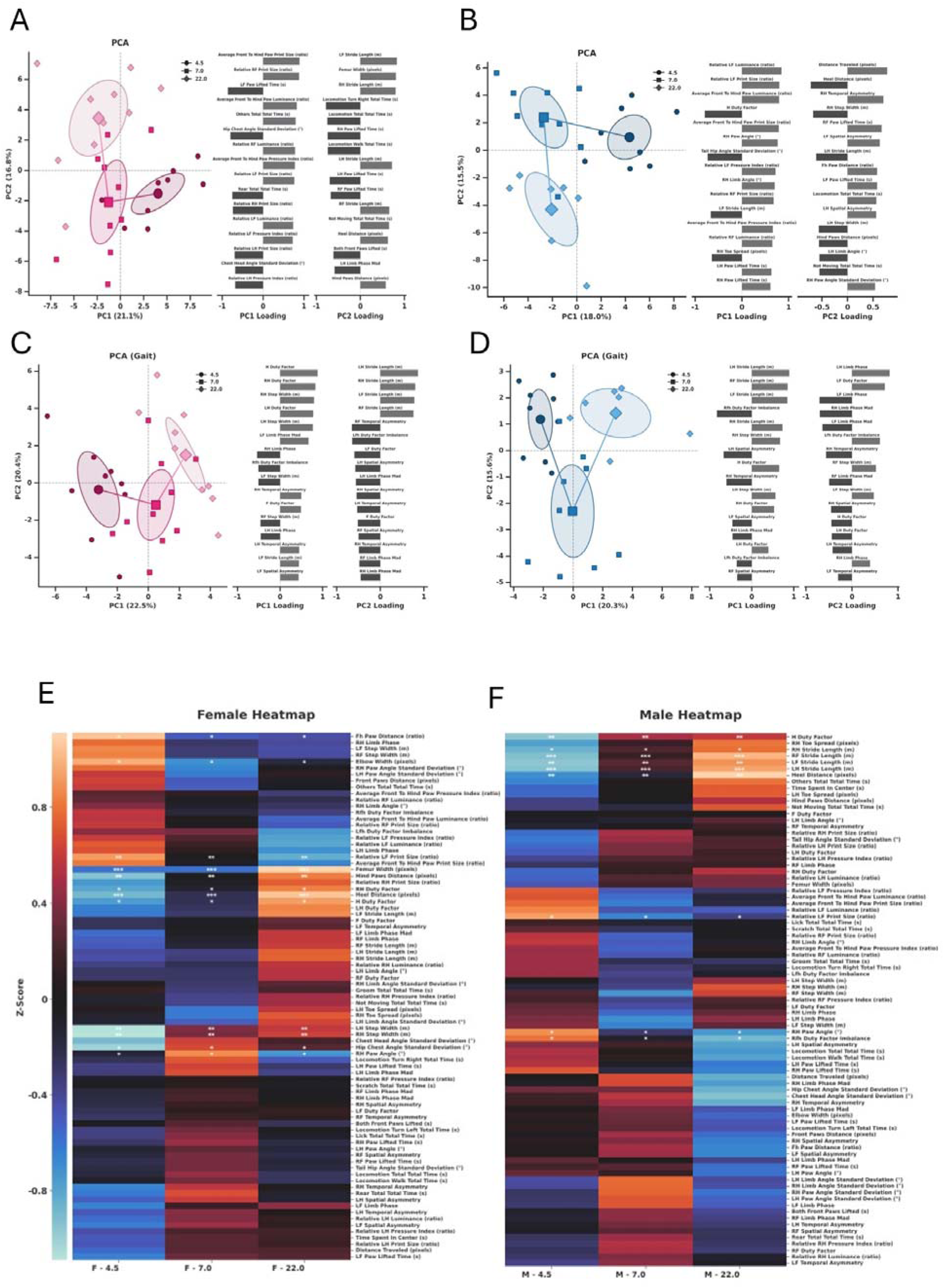
Age-related differences in multivariate spontaneous movement and gait phenotypes in male and female mice. Principal component analysis (PCA) of BlackBox-derived movement parameters from C57BL/6 mice at 4.5, 7, and 22 months of age. Each point represents one mouse (56 observations from 56 mice). (A-B) PCA of 85 BlackBox-derived features in females (A) and males (B), including measures of locomotor activity, gait, paw placement and loading, and posture. (C-D) PCA restricted to 34 gait-related parameters in females (C) and males (D). Data were covariate adjusted by body mass, linear, and angular velocity where applicable, then standardized within each PCA plot. Model marginal estimated means originated from the full ordinary least-squares fit using sex and age as fixed effects. No subject-level random intercept was included because each mouse contributed a single observation. PCA was performed using model-adjusted values from the groups shown in the analysis. Side panels show the 16 features with the highest loadings for PC1 and PC2. Ellipses indicate the 95% confidence interval (chi-square quantile) of each age’s estimated marginal mean. (E-F) Exploratory heatmaps of female (E) and male (F) features (the 85 included in PCA) after covariate-adjustment and z-scoring across each heatmap individually. Rows are ordered by hierarchical clustering separately within each panel. Asterisks mark features that differ among the three age groups by one-way ANOVA within sex, with Bonferroni correction across included features and repeated across the row (* p < 0.05, ** p < 0.01, *** p < 0.001).

When analyses were restricted to gait-related measurements, the age-related pattern became more distinct (**Figure 1C-D**). In females, the separation of the age groups was driven by coordinated differences in gait characteristics, including duty factor (a measure of the proportion of the gait cycle that a paw remains in contact with the ground), stride length and width, paw placement, and measures of limb asymmetry. The 7-month-old females generally showed an intermediate gait phenotype, while the 22-month-old mice showed a more distinct shift from the young group. The aged males showed a clear shift in gait phenotype compared with the younger groups. As in females, this separation reflected coordinated changes across several gait features, including duty factor, stride characteristics, paw placement, and limb asymmetry, rather than a change in a single gait measurement.

Exploratory heatmaps (**Figure 1E-F**) were used to further describe features included in PCA (**Figure 1A-B)**. These analyses more directly showed how groups of features were altered together thanks to hierarchical clustering of rows within each sex’s plot. For example, in males many paw-related angle standard deviations tended to be decreased all together at 22 months as compared to 7 months of age, even when the Bonferroni-corrected ANOVA did not identify them as statistically significant differences. Taken together, heatmap-based analysis of variables highlighted endpoints trends that may have been compressed by PCA, and effectively reflected all changes observed between groups rather than just those with the largest between-group differences.

### Aging differentially affects individual components of spontaneous locomotor activity

Individual measures of spontaneous locomotor activity showed different patterns with age and between sexes (**Figure 2**). In females, distance traveled and locomotion/walk time did not significantly differ among the 4.5-, 7-, and 22-month-old groups (**Figure 2A, C**), although variability in several measures appeared greatest in the older mice. In males, distance traveled was significantly lower in 22-month-old mice as compared to the 7-month-old mice (**Figure 2B**; p<0.05). Locomotion/walk time also tended to decrease with age, with 22-month-old males spending less time walking than 4.5-month-old males (**Figure 2D**; trend, p < 0.1).

**Figure 2.**
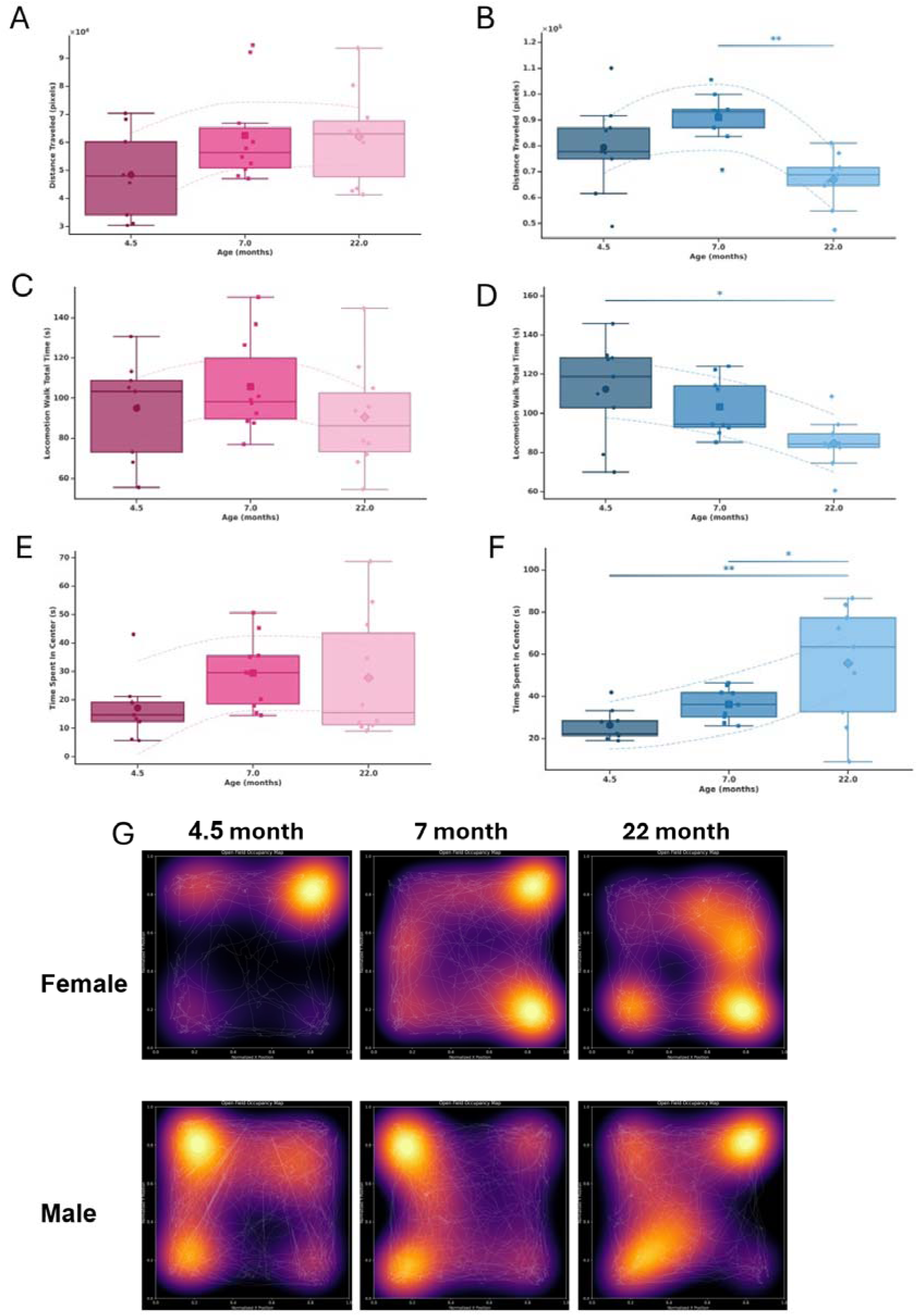
Age-related changes in spontaneous locomotor activity in female and male mice. BlackBox-derived measures of spontaneous activity in C57BL/6 mice at 4.5, 7, and 22 months of age. (A-B) Total distance traveled in females (A) and males (B). (C-D) Total locomotion/walk time in females (C) and males (D). (E-F) Time spent in the center in females (E) and males (F). Data were analyzed using ordinary least squares models with sex and age as fixed effects. Body mass was included as a z-scored covariate for distance traveled and time spent in the center. Model-based estimated marginal means and 95% confidence intervals are shown. Pairwise age-group comparisons were performed using model-based contrasts with Benjamini-Hochberg false discovery rate (FDR) correction for multiple comparisons. Individual points represent observations from each mouse. *p < 0.1, **p < 0.05, ***p < 0.001.

Time spent in the center of the analysis field showed a different age-related pattern (**Figure 2E-F**). While no significant differences were detected between groups in females, 22-month-old males spent more time in the center of the field as compared with both 4.5- and 7-month-old males (**Figure 2F**; p < 0.05 and p < 0.1, respectively). Notably, the aged male group also showed greater variability in center time as compared to the younger groups (**Figure 2G**).

### Aging is associated with coordinated changes in gait mechanics

Consistent with the gait-specific PCA, several individual gait parameters changed with age (**Figure 3**). Hindlimb duty factor increased with age in both females and males, indicating that older mice spent a greater proportion of the gait cycle with the hind paw in contact with the surface. In females, hindlimb duty factor was greater at 22 months of age than at both 4.5 and 7 months of age (**Figure 3A**; p < 0.05 for both comparisons). In males, this change was already present by 7 months of age, with both 7- and 22-month-old mice showing greater hindlimb duty factor than 4.5-month-old mice **(Figure 3B**; p < 0.001).

**Figure 3.**
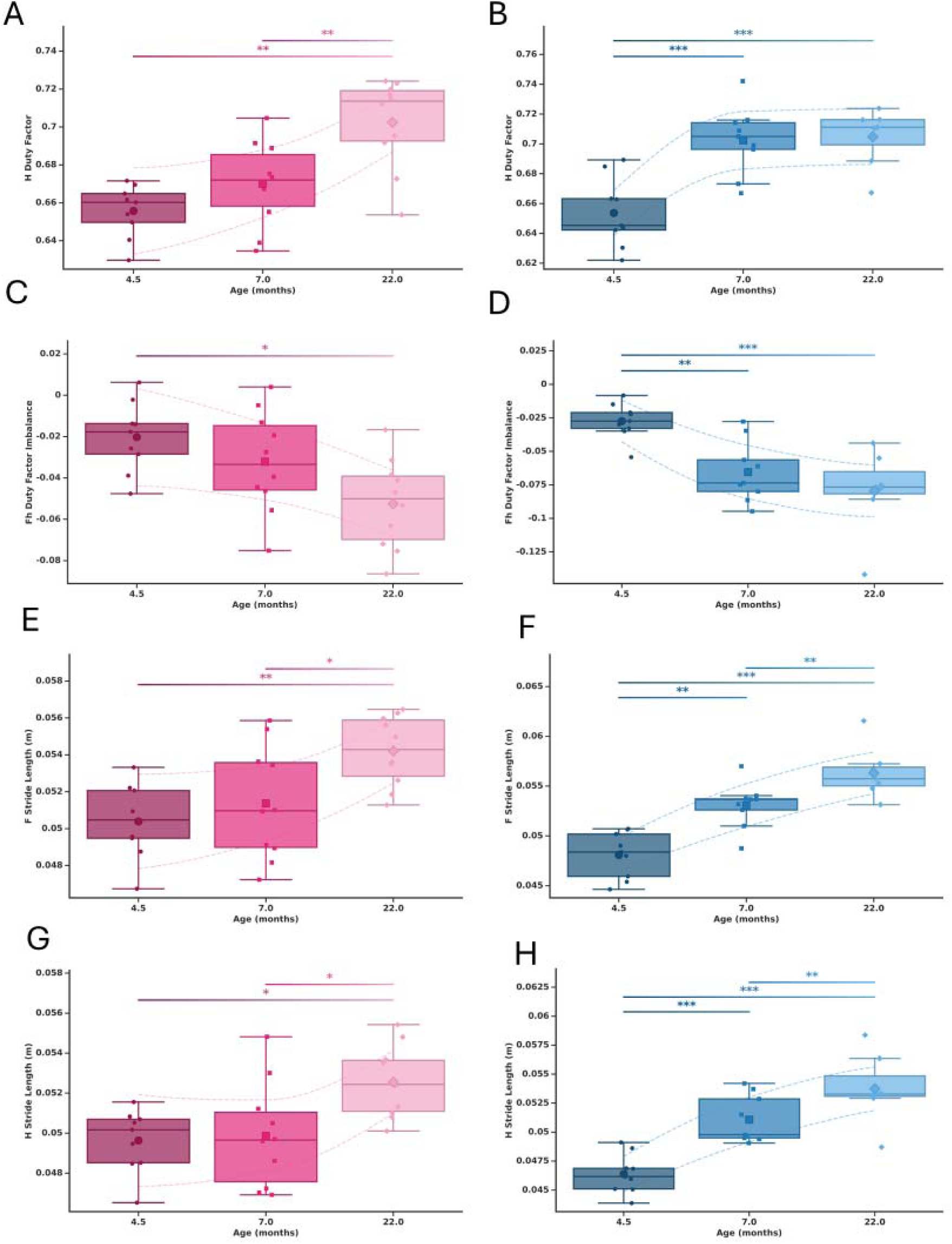
Age-related changes in gait mechanics in female and male mice. BlackBox-derived gait measurements in C57BL/6 mice at 4.5, 7, and 22 months of age (54 mice, 2 had less than 6 gait bouts and were excluded). (A-B) Hindlimb (H) duty factor in females (A) and males (B). (C-D) Fore to hind duty-factor imbalance in females (C) and males (D), calculated as the difference of the mean fore and hind duty-factors. (E-F) Forelimb (F) stride length in females (E) and males (F), calculated as the mean of left and right forepaw stride length. (G-H) Hindlimb stride length in females (G) and males (H), calculated as the mean of left and right hind paw stride length. Data were analyzed using ordinary least-squares models with sex and age as fixed effects and angular velocity, body mass, and velocity included as z-scored covariates. Each mouse contributed a single observation; therefore, no subject-level random effect was included. Model-based estimated marginal means and 95% confidence intervals are shown. Pairwise age-group comparisons were obtained from the fitted models and adjusted for multiple comparisons using the Benjamini-Hochberg false discovery rate (FDR) procedure. Individual points represent individual mice. *p < 0.1, **p < 0.05, ***p < 0.001.

Fore to hind duty-factor imbalance also changed with age (**Figure 3C-D**). Females showed a trend toward greater imbalance at 22 months compared with 4.5 months of age (**Figure 3C**; p < 0.1), indicating the hind paws had a greater ratio of contact time to stride time. In males, the difference was already apparent at 7 months of age as compared to 4.5 months of age and remained present at 22 months of age (**Figure 3D**; p < 0.05 and p < 0.001 vs. 4.5 months, respectively). Thus, aging was associated not only with longer relative hindlimb contact time, but also with changes in the balance of contact time between the forelimbs and the hindlimbs.

Stride length also increased with age when controlling for velocity (**Figure 3E-H**). In females, forelimb stride length was greater at 22 months of age than at 4.5 months of age (p < 0.05), with a trend toward an increase compared with 7 months (p < 0.1; **Figure 3E**). Hindlimb stride length showed a similar pattern, with trends toward longer strides at 22 months compared with both younger groups (p < 0.1; **Figure 3G**). The age-related increase in stride length was more pronounced in males. Forelimb stride length in males was greater at 7 months than at 4.5 months of age (p < 0.05) and was greater still at 22 months of age (p < 0.001 vs. 4.5 months; p < 0.05 vs. 7 months; **Figure 3F**). A similar pattern was observed for hindlimb stride length, which was greater at 7 months (p < 0.001 vs. 4.5 months) and remained greater at 22 months (p < 0.001 vs. 4.5 months; p < 0.05 vs. 7 months; **Figure 3H**) as compared to the youngest animals.

Together, these findings identify an age-associated gait phenotype characterized by increased hindlimb duty factor, altered fore to hind duty-factor balance, and longer strides. These changes were already apparent by 7 months of age in males, whereas in females they were more prominent at 22 months of age, suggesting that the timing and magnitude of age-related changes in gait may differ between sexes.

### Aging alters paw contact and limb-loading patterns

Age-related differences extended beyond stride and duty factor to measures of paw contact and paw-lift timing throughout the whole trial (**Figure 4**). The front-to-hind paw-print size ratio decreased with age after adjustment for body mass, indicating a change in the relative paw contact area between the forelimbs and hindlimbs. In females, the ratio was lower at 7 months than at 4.5 months of age (**Figure 4A**; p < 0.1) and was further reduced at 22 months of age (**Figure 4A**; p < 0.05 vs. 4.5 months). In males, the ratio was significantly lower at 7 months than at 4.5 months (**Figure 4B**; p < 0.05), while the 22-month-old group showed an intermediate phenotype. Paw-lift timing also changed with age, particularly in males (**Figure 4C-D**). Hind-paw lift time did not significantly differ among age groups in females (**Figure 4C**). In males, however, hind-paw lift time progressively decreased with age, with 22-month-old mice showing a shorter lift time than 4.5-month-old mice (**Figure 4D**; p < 0.05).

**Figure 4.**
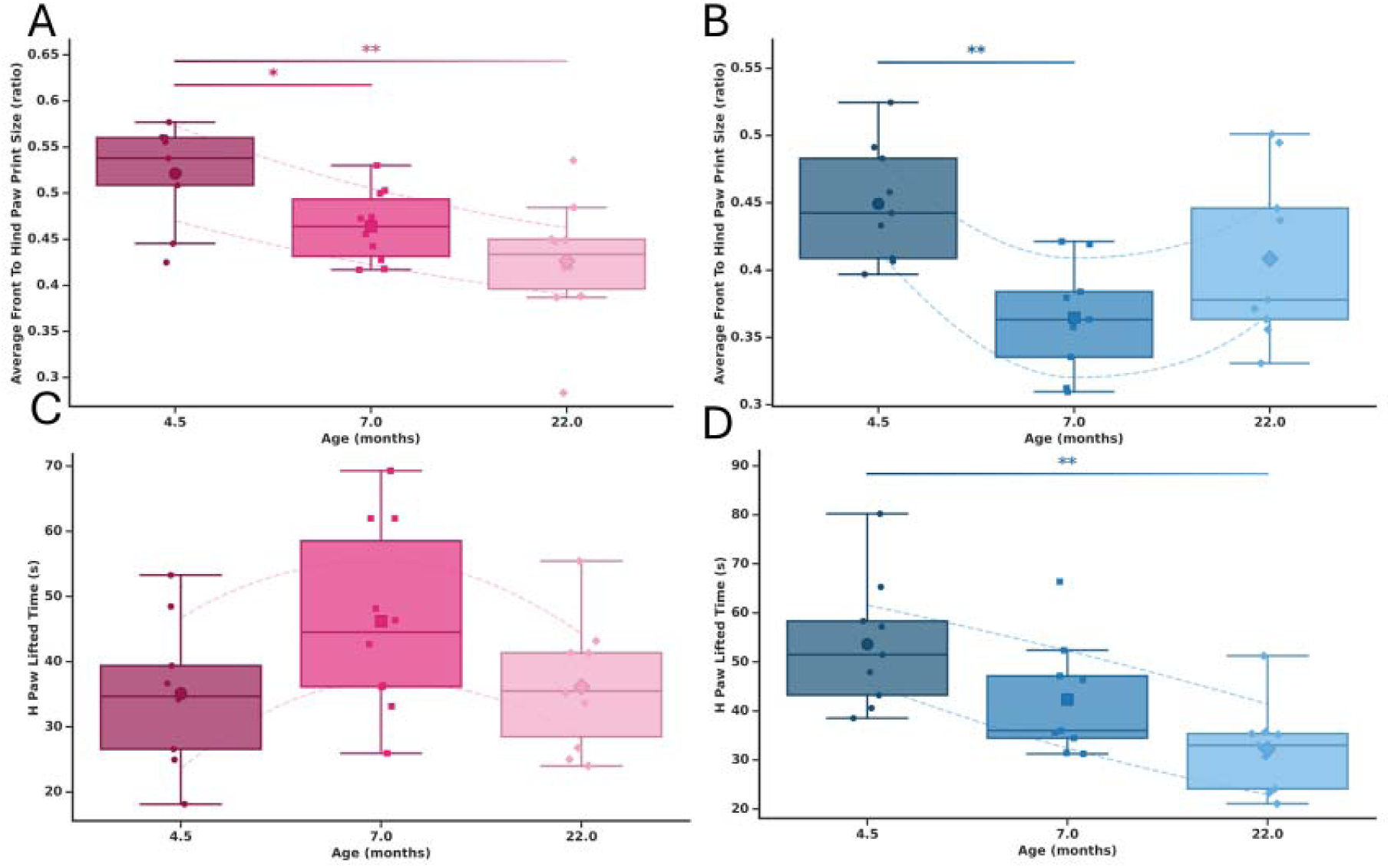
Age-related changes in paw contact and paw-lift timing in female and male mice. BlackBox measurement of paw contact and movement timing in C57BL/6 mice at 4.5, 7, and 22 months of age (56 mice). (A-B) Average front-to-hind paw-print size ratio in females (A) and males (B). (C-D) Hind-paw lifted time in females (C) and males (D), calculated as the mean of the left and right hind-paw lifted times. Data were analyzed using ordinary least-squares models with sex and age as fixed effects and body mass included as a z-scored covariate. Each mouse contributed a single observation; therefore, no subject-level random effect was included. Model-based estimated marginal means and 95% confidence intervals are shown. Pairwise age-group comparisons were obtained from the fitted models and adjusted for multiple comparisons using the Benjamini-Hochberg false discovery rate (FDR) procedure. Individual points represent individual mice. *p < 0.1, **p < 0.05, ***p < 0.001.

### Postural organization and variability change with age

Age-related differences were also observed in body position and postural organization during spontaneous movement (**Figure 5**). Front-paw distance decreased with age in both sexes. In females, front-paw distance was lower at 7 months than at 4.5 months (**Figure 5A**; p < 0.1). In males, front-paw distance was lower at 22 months than at 7 months (**Figure 5B**; p < 0.1). Heel distance increased with age. In females, heel distance was greater at 7 and 22 months compared with 4.5 months (**Figure 5C**; p < 0.05 and p < 0.001, respectively) and was further increased at 22 months compared with 7 months (**Figure 5C**; p < 0.001). A similar pattern was observed in males, with greater heel distance at 22 months compared with both 4.5 and 7 months (**Figure 5D**; p < 0.001 and p < 0.05, respectively).

**Figure 5.**
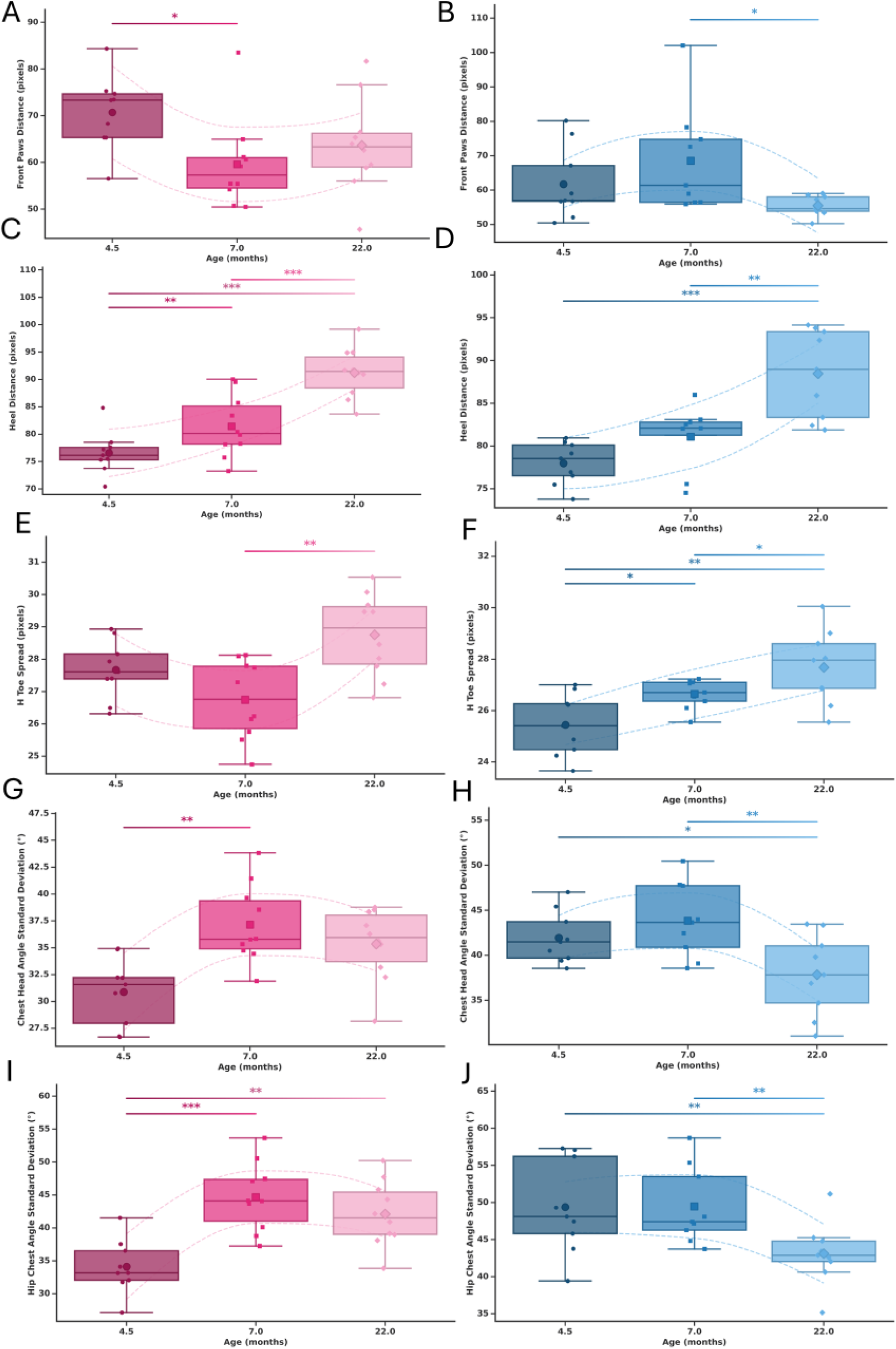
Age-related changes in postural organization and variability in female and male mice. BlackBox-derived parameters of body geometry and postural variability in C57BL/6 mice at 4.5, 7, and 22 months of age. (A-B) Front-paw distance in females (A) and males (B). (C-D) Heel distance in females (C) and males (D). (E-F) Hind-toe spread in females (E) and males (F). (G-H) Variability in chest-head angle, expressed as the standard deviation (SD), in females (G) and males (H). (I-J) Variability in hip-chest angle, expressed as the SD, in females (I) and males (J). Individual points represent individual mice. Box plots show the distribution within each age group, and dashed lines connect group model marginal estimated means’ 95% confidence intervals to illustrate age-related trends. Pairwise age-group comparisons were adjusted for multiple comparisons using the Benjamini-Hochberg false discovery rate (FDR) procedure. *p < 0.1, **p < 0.05, ***p < 0.001.

Hind-toe spread also changed with age when adjusted for body mass (**Figure 5E-F**). In females, hip-toe spread was greater at 22 months than at 7 months **(Figure 5E**; p < 0.05). In males, hind-toe spread increased at 7 months compared with 4.5 months (**Figure 5F**; p < 0.1) and was greater at 22 months than at 4.5 months (p < 0.05), with a trend toward a further increase between 7 and 22 months (**Figure 5F**; p < 0.1).

Age also affected the variability of body position. In females, variability in the chest-head angle was greater at 7 months than at 4.5 months (**Figure 5G**; p < 0.05). Hip-chest angle variability was also greater at both 7 and 22 months compared with 4.5 months (**Figure 5I**; p < 0.001 and p < 0.05, respectively). In males, however, the pattern differed. Chest-head angle variability was lower at 22 months than at 7 months (**Figure 5H**; p < 0.05), while hip-chest angle variability was lower at 22 months than at 4.5 months (**Figure 5J**; p < 0.05).

## DISCUSSION

The current study suggests that aging can alter patterns of movement (“<u>how</u> mice moved”) more consistently than the overall amount of movement (“how <u>much</u> they moved”). Gait-specific principal component analysis showed age-related separation between the 4.5-, 7- and 22-month-old groups, supported by changes in hindlimb duty factor, stride length, duty-factor imbalance, paw contact, and posture. In contrast, total distance traveled and walking time showed fewer age-related changes and more sex-dependent differences. These findings suggest that gait and postural measures may detect age-related functional changes that are missed when spontaneous activity is assessed only by distance or time spent moving. Most importantly, the PCA- and heatmap-based visualization of results permitted simultaneous consideration of 85 of all 200+ variables tracked by the Palmreader software (and engineered features) in an efficient manner, permitting identification of age-related trends in nuanced aspects of gait and posture that may have been otherwise missed by conventional approaches used to track physical activity.

The age-related changes in voluntary activity metrics identified here are relevant to estimates of frailty and intrinsic capacity because commonly used measurements to characterize these categorial phenotypes include walking speed, strength, endurance, body weight, and physical activity [26]. Frailty measured using these approaches increases with age and is associated with mortality in C57BL/6 mice [25]. The newer concept of intrinsic capacity considers locomotor capacity as a key contributing variable to healthy aging, which in animal models encompasses measurements of endurance, balance, and muscle strength such as wheel running, rotarod performance, and treadmill testing [11]. The BlackBox One system studied here may be capable of complementing these measures by providing information about how movement is performed, including duty factor, fore-hind shifts, stride characteristics, and postural variability. However, the measures identified here should currently be considered age-sensitive movement measures rather than validated frailty or locomotor capacity markers, particularly because some differences were already present at 7 months as compared to 4.5 months of age, well before mice would typically be considered frail. Despite this caveat, the noted advantage of the current approach is the ability to generate a wide scope of locomotor-related metrics that demonstrate age-related functional changes in locomotor activity in a short timeframe (7-minute data recording period per animal following a single group-based acclimation period) with little to no direct handling of the animals involved. This approach mitigates concerns related to sources of stress and inter-operator variability, especially in aged animals, that may advance understanding of functional musculoskeletal decline during aging.

It is important to note that some of our findings differ from previous reports. For example, the aged mice in our study demonstrated longer forelimb and hindlimb strides as compared to younger groups, whereas an earlier study using freely moving male C57BL/6 mice reported decreased cadence and increased stride-time variability with aging but no significant change in stride length or duty cycle [42]. This discrepancy may be due to different ranges of included velocities, number of bouts (in our study the minimum number of bouts recorded was 24, and the average closer to 100) and covariate adjustment methods. Our velocity adjusted increase in stride length corresponds with decreased cadence reported previously [42]; indeed, further investigation of stride length divided by velocity showed significant increase in stride time (*not shown*). Other work has shown that C57BL/6 mice can develop longer and less frequent strides with age, mainly during earlier adulthood [43] which is more directly consistent with our findings. Longer strides in our aged mice therefore should not be interpreted as improved function; aged males traveled less while showing longer strides and greater hindlimb duty factor, suggesting a different movement strategy with age rather than having superior locomotor performance.

Some limitations of our study should be considered. In addition to the fact that the youngest animals (4.5 months of age) were sourced from in-house breeding rather than the parent colonies of the 7-month-old and 22-month-old mice, this study was also cross-sectional rather than longitudinal in nature and included a relatively modest number of animals in each group. We also have not yet directly compared the metrics quantified in this study with established frailty indices, strength, endurance, or survival. Therefore, it remains unknown whether these movement changes predict frailty or simply reflect patterns of normal aging, which will be more thoroughly interrogated in future studies. Longitudinal studies combining BlackBox-facilitated measurements with established frailty and functional assessments will be needed in the future.

Overall, our study identified age-related changes in gait, paw contact, and posture of male and female mice that are not fully captured by conventional activity measures. The potential value of this approach, in the context of aging-related musculoskeletal decline, is the ability to measure multiple components of spontaneous movement, behavior, posture, and weightbearing together and repeatedly across a single cohort of animals. Future studies could determine whether a combination of these measures can detect the transition from normal aging to frailty and provide a practical tool for longitudinal studies of physical decline.

## Acknowledgements

This study was supported by the NIH (P01 AG036675 Core B and Project 4, U01 AG086158, R01 AR082307). Funding from the American Association for Laboratory Animal Science supported the generation of the 4.5-month-old animals that were studied.

